# RISC: robust integration of single-cell RNA-seq datasets with different extents of cell cluster overlap

**DOI:** 10.1101/483297

**Authors:** Yang Liu, Tao Wang, Deyou Zheng

## Abstract

Single cell RNA-seq (scRNA-seq) has remarkably advanced our understanding of cellular heterogeneity and dynamics in tissue development, diseases, and cancers. Integrated data analysis can often uncover molecular and cellular links among individual datasets and thus provide new biological insights, such as developmental relationship. Due to differences in experimental platforms and biological sample batches, the integration of multiple scRNA-seq datasets is challenging. To address this, we developed a novel computational method for robust integration of <u>sc</u>RNA-seq (RISC) datasets using principal component regression (PCR). Because of the natural compatibility of eigenvectors between PCR model and dimension reduction, RISC can accurately integrate scRNA-seq datasets and avoid over-integration. Compared to existing software, RISC shows particular improvement in integrating datasets that contain cells of the same types (more accurately clusters) but at distinct functional states. To demonstrate the value of RISC in finding small groups of cells common between otherwise heterogenous datasets, we applied it to scRNA-seq datasets of normal and malignant cells and successfully identified small clusters of cells in healthy kidney tissues that may be related to the origin of renal tumors.

## Introduction

Single-cell RNA-seq has become an essential genomic technology (Islam et al. 2014; Nawy 2014; Wang and Navin 2015; Zheng et al. 2017) and it is now widely used in many biological domains. Such domains include surveying cell heterogeneity among embryonic stem cells and tumors (Azizi et al. 2018; Rosenberg et al. 2018), identifying genetic markers of specific cell types (Fan et al. 2018), and investigating cell fate commitment and lineage trajectories (Wang et al. 2017). As shown in a recent study describing the lineages and trajectories of aging Drosophila brain cells, the integration of scRNA-seq datasets from different studies can provide a high level understanding of cellular heterogeneity and functional relationship across various tissue types or development stages (Davie et al. 2018). However, noise from experimental batches, assay conditions, sequencing platforms, and other biological and technical factors have made the task extremely challenging. Due to the unique nature of scRNA-seq data (e.g. low coverage and dropout) integration methods designed for bulk RNA-seq data are usually not applicable to scRNA-seq analysis. In addition, existing scRNA-seq analytical software typically normalizes scRNA-seq data by scaling raw counts or unique molecular identifiers (UMIs) to account for the difference in sequencing depth, ignoring more complicated heterogeneity between different datasets.

Two approaches have recently been developed to integrate scRNA-seq datasets. One is embedded in an R toolkit for single cell genomics called “Seurat,” which uses Canonical Correlation Analysis (CCA) (Butler et al. 2018). The other is “Scran” that applies the Mutual Nearest Neighbors (MNN) algorithm (Haghverdi et al. 2018). Seurat utilizes principal CCA to generate a covariance matrix from gene expression matrices of different studies, decomposes the covariance matrix by singular-value decomposition (SVD), generates eigenvectors, and embeds the cells based on the eigenvectors. Similar to Seurat, Scran performs MNN to define cell pairwise relationships by correcting gene expression matrices before CCA.

These approaches are valuable for integrating scRNA-seq datasets with relatively homogeneous cell types (Butler et al. 2018; Haghverdi et al. 2018). How to integrate datasets with high-level of heterogeneity in cell types remains a challenge, for example, in the study of developmental lineages with a commonality only in their progenitor cells. Furthermore, it remains to be addressed whether an integration software can distinguish cells of the same type but at different functional states. Therefore, we developed a new toolkit, RISC, with an aim to integrate scRNA-seq datasets with both high and low level of cell cluster similarity, which is defined herein as the proportion of shared cell types (or clusters) between datasets, because the cell population diversity of a sample is determined more by the number of cell clusters than by the numbers of cells in each cluster. The main component of our approach utilizes principal component regression (PCR) model, which effectively embeds single cells by the eigenvectors derived from the gene expression matrix of individual datasets and uses these eigenvectors to integrate datasets (Jolliffe 1982; Bair et al. 2006).

We evaluated the performance of RISC with multiple scRNA-seq datasets containing various degrees of cell cluster similarities (Hashimshony et al. 2012; Jaitin et al. 2014; Picelli et al. 2014; Zheng et al. 2017), and also compared it to Seurat and Scran. These datasets were all from previous studies in very distinct biological contents, ranging from blood cells, pancreas cells, immune cells in breast tumor microenvironment, to kidney tumors. They were generated by multiple platforms, including cell expression by linear amplification and sequencing (CEL-seq), massively parallel single-cell RNA-sequencing (MARS-seq), Full-length RNA-seq (SMART-seq2), and 10x Genomics’ Chromium system (Hashimshony et al. 2012; Jaitin et al. 2014; Picelli et al. 2014; Zheng et al. 2017). The results indicate that RISC can successfully merge cells sharing common gene expression signature, while accurately preserving the separation of non- intersected cells from different datasets.

## Results

### Overview of RISC

Dimension reduction is the fundamental step in scRNA-seq analysis because cell embedding, cell clustering, cluster marker identification, and all other subsequent analyses depend on its result. It often relies on principal eigenvectors to represent the high-dimensional data (Roweis and Saul 2000a). Since the eigenvectors of dimension reduction capture most biological information of individual dataset, they can be effectively exploited for data integration. Because of the lack of selection and regularization, algorithms like CCA directly combine all the information of gene expression matrices across datasets including the noises that have been discarded in the first step of dimension reduction of individual datasets (Hotelling 1933; Roweis and Saul 2000b; Pierson and Yau 2015; Ding et al. 2018). This noise can contribute to under or over integration. To address this, RISC utilizes PCR model to align the principal components (PCs) from the dimension reduction (Hotelling 1936). As such, the subsequent analyses of the integrated data will be maximally consistent with the dimension reduction results of individual datasets (see Methods).

Additionally, both dimension reduction and data integration in scRNA-seq datasets are mainly based on the genes with the highly variable expression. As a result, an input matrix for integration has the number of genes (rows) similar to or less than the number of cells (columns). This kind of matrices cannot be decomposed to generate unique SVD (Cheney and Kincaid 2007) and one cannot directly perform CCA on them without appropriate regularization (Witten et al. 2009). To handle this problem, RISC utilizes a subset of PCs for regression model and thus forming an effective regularized procedure (see Methods) (Bair et al. 2006).

Here, we briefly explain the key steps of the algorithm and features of RISC (Fig. 1A; details in Methods).

**Figure 1.**
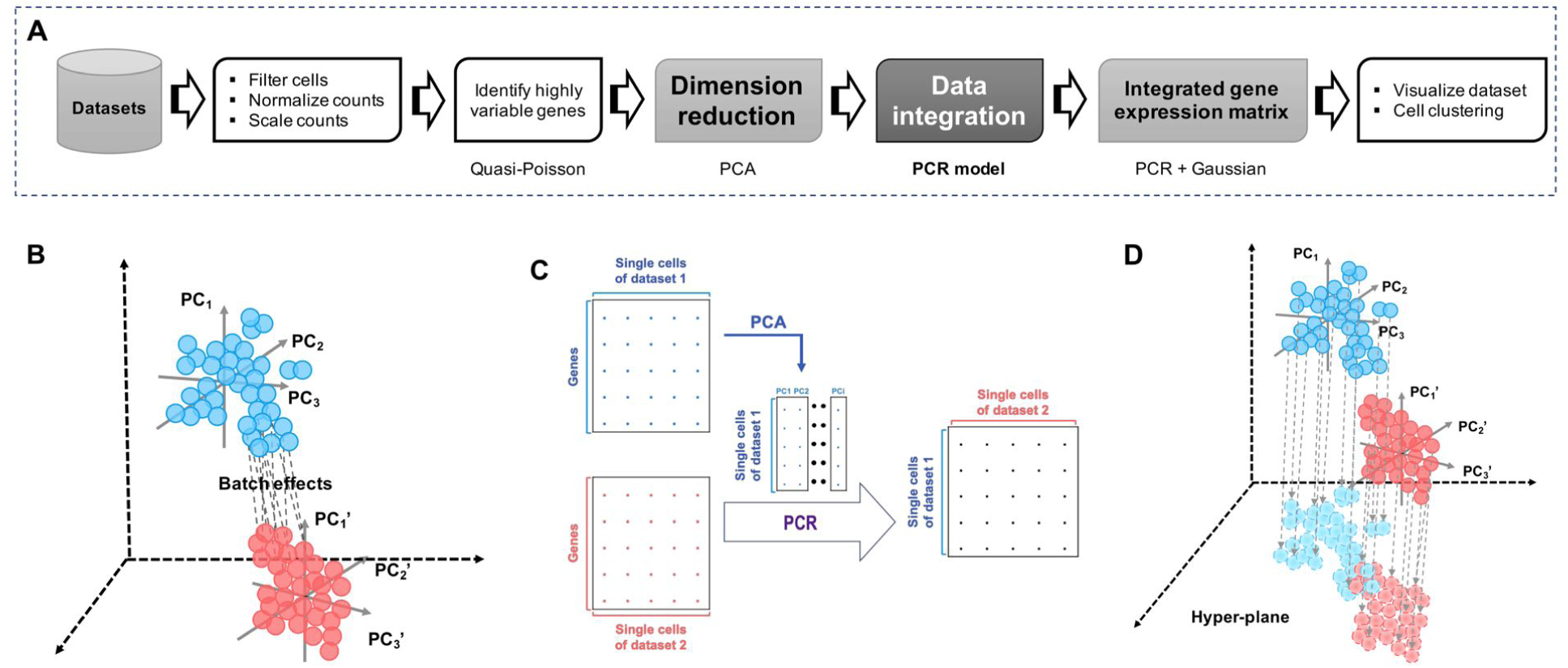
The schematic workflow of RISC. (A) Flowchart of the main steps in RISC. (B) Difference in dimension space between two scRNA-seq datasets represented as batch effects, with dash lines connecting cells of the same type in different datasets. (C) RISC transfers high-dimension gene expression matrices into PCs and utilizes these PCs to correct batches by PCR model and generate an integrated dataset. (D) Two scRNA-seq datasets are integrated in a hyper-plane with covariance in appropriate fit-error between them.

1. RISC pre-processes the gene expression matrix in each dataset to make it ready for integration. For simplicity of description, we consider the global difference between any two datasets as “batch effects” (Fig. 1B), which represent technical noises in most cases (Leek et al. 2010) but can be biological differences. First, RISC normalizes the gene expression values of each cell by a size factor that is derived from the total count/UMI in that cell. Next, it scales the gene matrix to balance expression levels in each cell with empirical mean equal to 0. Lastly, it identifies genes exhibiting highly variable expression via a Quasi-Poisson model.
2. In the process of batch correction and integration (Fig. 1C), RISC first transfers the expression matrix for the highly variable genes in the reference dataset into PCs. The default in RISC chooses the dataset with the largest cell number as the reference, but a user can modify as they see fit. The PCs chosen for PCR-integration are from the PCs (either all or a subset) used for dimension reduction of individual datasets. This novel strategy of RISC avoids loss of information or introduction of new noise by integration itself. Then, RISC derives a covariance hyper-plane between the reference and target datasets using the PCR model (Fig. 1D). Lastly, RISC adjusts gene expression ranges of each gene across datasets to generate an integrated gene expression matrix.
3. RISC also provides cell clustering function and methods for identifying cluster marker genes, using the previously described algorithm (see Methods). In addition to a typical choice of negative binomial regression, RISC implements a second option for computing cluster marker genes based on Quasi-Poisson regression that can stably detect marker genes in small numbers of cells (Ver Hoef and Boveng 2007).

### The general performance of RISC in data integration

To evaluate the data integration ability of RISC, we first examined its accuracy in merging cells belong to the same cell types. This was accomplished by the analysis of two publicly available scRNA-seq datasets of human peripheral blood mononuclear cells (PBMCs) (Kang et al. 2017), with raw data obtained from the Gene Expression Omnibus (GEO; accessible number GSE96583). One set was from PBMCs after interferon-β (IFN-β) treatment for 6 hours, while the other was controls. We chose to test RISC with these realistic datasets over computationally simulated ones because simulation would not be able to mimic all the known and unknown factors in scRNA-seq data production. Analysis of the raw dataset (termed “pre-integrated data sets” herein) by t-SNE (t-distributed stochastic neighbor embedding) showed a clear separation of the IFN+ cells and IFN- controls (Fig. 2A), which is most likely caused by batch effects, because Kang et al reported only a few hundreds of differentially expressed genes (DEGs) for most cell types upon IFN-β treatment (Kang et al. 2017). After RISC integration, the batch difference was removed as the separation of the two datasets disappeared (Fig. 2B). The necessary of batch correction for these two datasets was also described previously (Butler et al. 2018).

**Figure 2.**
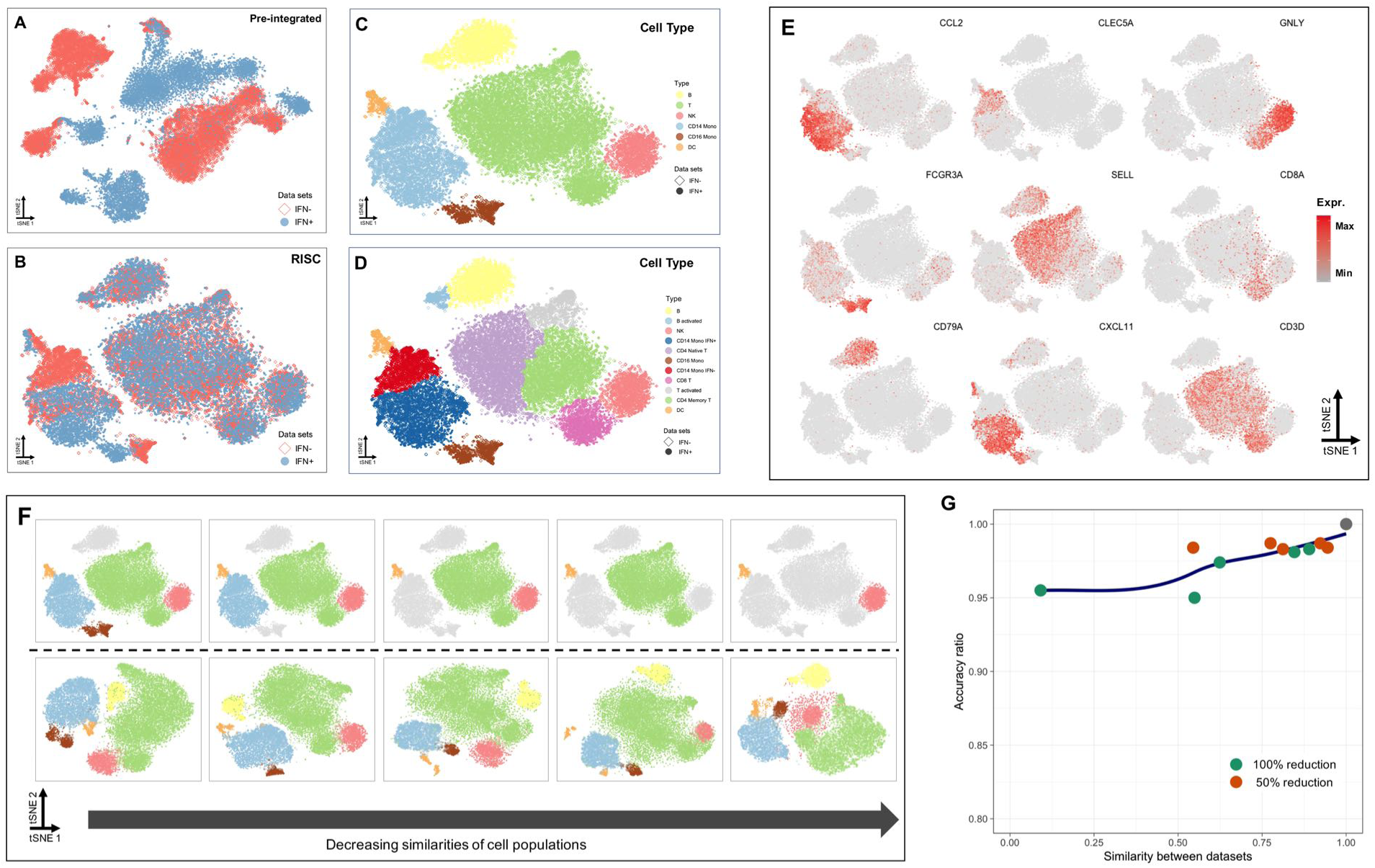
Performance of RISC in integration of PBMC scRNA-seq datasets with different cell composition similarities. (A ~ B) T-SNE plots display cell embedding of PBMC cells from control (“IFN-”) and IFN-β stimulation (“IFN+”), before (A) and after integration by RISC (B), with light red for “IFN-” data and light blue for “IFN+” data. (C ~ D) In t-SNE plots, the embedding cells of the RISC-integrated dataset are colored by cell types (C) or cell sub-populations (D), as defined previously (Kang et al. 2017). (E) The expression patterns of cell-type markers, corresponding to cell populations in D. (F) The upper plots indicate the generation of simulated data subsets with reducing cell cluster similarity (from left to right), in which the gray color marks the cells gradually removed from the full “IFN+” data. The lower plots show cell embedding of the integrated dataset between the “IFN-” data (full cell populations) and individual simulated “IFN+” data subsets. The colors for cell populations correspond to the panel C. (G) The accuracy of RISC in data integration with different cell clustering similarities, where the accuracy is estimated by the ratio of correct cell embedding.

To evaluate the accuracy of RISC, we applied clustering analysis to the integrated data and annotated the identities of cell clusters by known marker genes. Using the previously defined cell types (or clusters) as the reference standard, both by Seurat (Butler et al. 2018) and by Kang et al (Kang et al. 2017), RISC correctly merged cells belonging to the same types, including B cells, natural killer (NK) cells, CD16 monocytes (CD16 Mono), CD14 monocytes (CD14 Mono), Dendritic cells (DC), and T cells (Fig. 2C). T cells were further divided into sub-types, CD4 Naïve T cells, CD8 T cells, activated T cells, and CD4 memory T cells (Fig. 2D), supported by the expression patterns of corresponding gene markers (Fig. 2E). Interestingly, RISC divided the CD14 monocytes into two sub-clusters, with the smaller one containing 1,816 control and 98 IFN+ cells while the larger one containing 1,237 control and 2,745 IFN+ cells. We thus referred to the former and the latter as “CD14 Mono IFN-” and “CD14 Mono IFN+” clusters, respectively (Fig. 2D). Differential expression analysis detected 2,894 DEGs (adjusted p-value < 0.05), of which 1,149 were expressed higher in the IFN+ cluster, including CXCL11 (Fig. 2E). 42% (n = 1,218) of these DEGs were also identified as differentially expressed in CD14 monocytes by Kang et al, indicating that the difference between the two CD14 monocyte (sub)clusters largely reflects the effects of IFN-β stimulation and thus represents distinct functional states. Note that this finding was not observed in a previous integration analysis (see below) (Butler et al. 2018). The significance of our result is further supported by the fact that more than two thirds of the total DEGs from IFN-β stimulated PBMCs were observed in the CD14 monocytes (Kang et al. 2017). The cell type with the second most DEGs were CD16 monocytes, which also seemed to contain two sub-clusters (Fig. 2B, D). Overall, these results indicate that RISC is not only able to integrate cells by types correctly but also preserve the treatment effect sufficiently.

Next, we investigated the RISC performance under various cell-population similarities between datasets, using the two PBMC datasets described above. We kept all cells in the “IFN-” dataset, but gradually removed the cells (either all or by 50% Fig. S1) of B, T, NK, CD14 Mono, CD16 Mono, and DC cells from the “IFN+” sample to generate data subsets with reduced similarities at the cell type level to the “IFN-” data (Fig. 2F top). Each of the modified “IFN+” data subsets were then integrated with the full “IFN-” data by RISC (Fig. 2F bottom). Then, we evaluated the accuracy of data integration by examining how the remaining cells in the “IFN+” data were clustered with the matching “IFN-” cell types. As shown in Fig. 2F and Fig. S1A, RISC’s performance was robust to cell similarity changes as it consistently aligned >90% of the “IFN+” cells to the correct cell types in the full “IFN-” data even when only ~10% of cell similarity existed between “IFN-” and “IFN+” datasets (Fig. 2G). In short, this analysis shows that RISC can accurately merge cells of the same types even when two data sets share very limited similarity, an important feature not addressed in previous integration software.

### Integration performance of RISC on datasets with highly comparable cell type composition

We then compared RISC’s performance to two existing software (Butler et al. 2018; Haghverdi et al. 2018) by analyzing multiple scRNA-seq datasets, starting from the data collected for pancreatic islets of two species, human and mice, which were consisted of nearly identical cell types (GEO: GSE84133) (Baron et al. 2016). Single cells from both species were captured in droplet microfluidics and sequencing libraries were prepared by the CEL-seq protocol (Hashimshony et al. 2012); therefore, the main factors to address in integration were species and donor differences (four human donors and two mouse donors). As shown by t-SNE plot, cells in the pre-integrated datasets were separated by species (Fig. 3A) and cell types (Fig. 3B). After integration by either RISC or Seurat, both species and donor differences disappeared and cells were separated by cell types, according to the cell type annotation provided by Baron et al (Baron et al. 2016). The cell embedding results confirmed the accuracy of the integrated datasets, as cell types mostly merged correctly, including acinar, alpha (α), beta (β), gamma (γ) delta (δ) cells, ductal, endothelial, stellate and macrophage cells (Fig. 3C, 3D, 3F and 3G). The performances of RISC and Seurat were similar, but the results from Scran, showed some degrees of mixture of the major cell types, alpha, beta, and delta cells (Fig. 3E, 3H).

**Figure 3.**
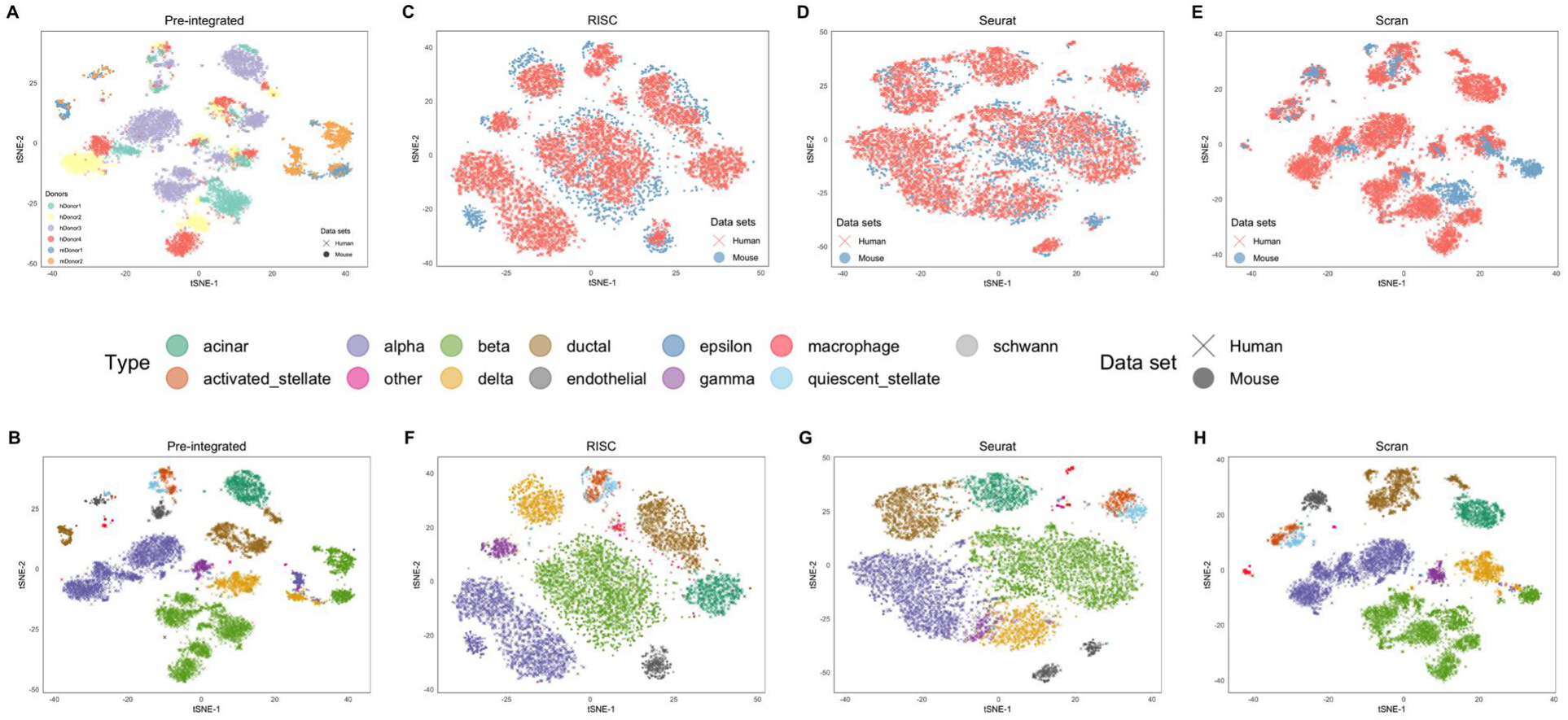
Integration of cross-species scRNA-seq datasets. (A) The t-SNE plot displays the pre- integrated datasets, six different colors representing different donors from human (n = 4) or mouse (n = 2). (B) According to the previous cell-type annotations (Baron et al. 2016; Butler et al. 2018), cells of the pre-integrated datasets are colored at t-SNE plot. (C ~ E) The integrated datasets by RISC (C), Seurat (D) and Scran (E), colored by species. (F ~ H) The integrated datasets by RISC (F), Seurat (G) and Scran (H), colored by cell types in B.

Next, we evaluated integration of datasets from different publications, which presumably have larger batch differences than datasets in the same study. We tested three pancreas scRNA-seq datasets (E-MTAB-5061, GSE81076 and GSE85241) with well-defined and annotated cell types (Grun et al. 2016; Muraro et al. 2016; Segerstolpe et al. 2016). The single cells of these sets were from healthy or type 2 diabetics (T2D) human donors, and three different protocols were used to obtain the scRNA-seq data, Smart-seq2 for E-MTAB-5061, CEL-seq for GSE81076, and CEL- seq2 for GSE85241. As expected, the cells were largely segregated by experimental protocols and donors before batch correction (Fig. S2A). After integration, RISC, Scran and Seurat all removed the batch effects and merged cells appropriately (Fig. S2B ~ D). Based on the original cell type annotation and the corresponding markers (Grun et al. 2016; Muraro et al. 2016; Segerstolpe et al. 2016), we examined the expression patterns of marker genes for four endocrine cell sub-populations (i.e., GCG, INS, PPY and SST for alpha (α), beta (β), gamma (γ) and delta (δ) cells, respectively). The results showed that all three methods successfully integrated data (Fig. S2E), but RISC and Seurat seemed to group cells of the same types more closely. Similar results were obtained when the three methods were used to integrate scRNA-seq datasets obtained by two different platforms for studying hematopoietic lineage (Smart-seq2 for GSE81682 and MARS-seq for GSE72857) (Paul et al. 2015; Nestorowa et al. 2016), as shown in Fig. S2F ~ J.

### Outperformance of RISC in preserving subtle subpopulation difference between datasets

A good integration software has to correctly align cells (from different datasets) that have the same gene expression profile, but it also needs to preserve the biological difference between datasets. In the integration of human PBMC datasets before and after IFN treatment, RISC was able to preserve the treatment effect on CD14 and CD16 monocytes (Fig. 2D), which was masked by Seurat (Fig. 4A ~ B and S3A ~ B). The RISC’s feature of avoiding over-integration is important for detecting cells that belong to the same cell type but are at distinct states, reflecting either developmental stages or in response to experimental treatments.

**Figure 4.**
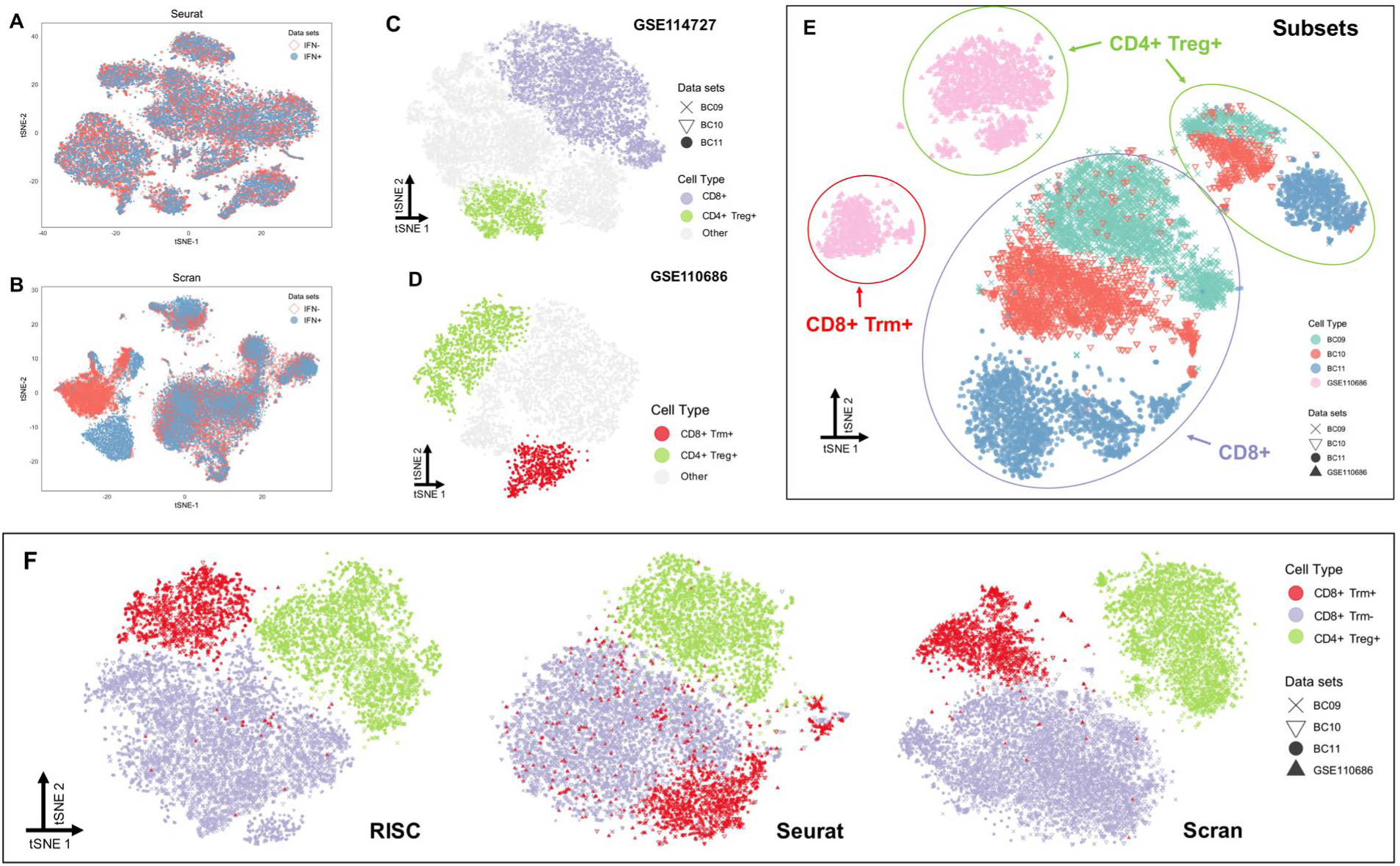
Separation of functionally distinct cells of the same type. (A ~ B) T-SNE plots show the integration results of the two PBMC datasets (Fig. 2) by Seurat (A) and Scran (B). (C) Separation of the CD8+ T and CD4+ Treg cells from other cells in the GSE114727 datasets. (D) Separation of the CD8+ Trm and CD4+ Treg cells from other cells in the GSE110686 dataset. (E) T-SNE plot shows the pre-integrated data for cells extracted from (C) and (D), with different cell types indicated by circles. (F) Integration of the four datasets in (D) by RISC (left), Seurat (middle) and Scran (right), with colors representing three types of T cells.

We then compared RISC, Seurat and Scran by analyzing immune cells in breast tumor microenvironment. The first datasets were obtained from the CD3+ leukocytes of three breast cancer patients using the 10X genomics’ protocol (GEO: GSE114727; “BC09”, “BC10” and “BC11”) (Azizi et al. 2018). Again, cells were separated by donor batch effects before integration (Fig. S3C), but RISC, Seurat and Scran mitigated the donor difference and correctly aligned cells of the same clusters (Fig. S3D ~ F). In the original study (Azizi et al. 2018), Azizi et al found that these CD3+ leukocytes could be divided into three groups (CD8+, CD4+ and regulatory T cells (Treg)), based on T cell activation signature genes (CD8A, CD4 and FOXP3). This was reproduced in the integrated datasets from RISC and Seurat, while some degree of under integration was seen in the Scran result (Fig. S3F, G). Nevertheless, the separation of the three cell types is somewhat unclear, suggesting a challenging dataset for integration and clustering.

To better illustrate the difference of the three software, we extracted the CD8+ and CD4+ Treg cells from the three donors (for better cluster separation) (Fig. 4C) and integrated them with the CD8+ T cells from an independent study that also investigated tumor-infiltrating T cells in breast cancer (Savas et al. 2018). From the second dataset (GSE110686), we extracted only CD4+ Treg cells and CD8+ tissue-resident memory T cells (CD8+ Trm) (Fig. 4D), which were shown to form a functionally distinct T cell group and provide better prognostication prediction than CD8+ T cells alone (Savas et al. 2018). Thus, Treg cells from the two reports were expected to be mixed but other cells (the majority) were not (Fig. 4E). To our surprise, after integration, we found that a small fraction of the CD8+ cells from the first report (i.e., GSE114727) actually showed characteristic gene expression of CD8+ Trm. This subtle but potential important finding was not described by Azizi et al, but was clearly supported by the expression of Trm cell markers (CCL3 and HAVCR2) (Savas et al. 2018) (Fig. S3H). We then checked the CD8+ Trm cells from the second report (i.e., GSE110686) and determined how many of them were mixed with non-Trm cells (i.e., Trm- CD8+) upon integration by RISC, Seurat and Scran (red cells in Fig. 4E and S3I). The result showed that RISC and Scran had good performance, with < 5% being aligned to the non-Trm CD8+ cluster. In contrast, >30% of the Trm+ cells were put incorrectly to the non-Trm cluster in the Seurat integrated data (Fig. 4F and S3I). Based on these results, we consider that RISC has a better performance in preserving subtle but biologically meaningful differences during data integration.

### Application of RISC for identifying tumor-like cells in normal human kidney tissues

As RISC can correctly identify and merge small clusters of common cells among samples (Fig. 2), we decided to test its capability in detecting tumor-like cells present in normal tissues, either next to tumors or before tumors emerge. In a recent study, Yang et al used a combination of scRNA-seq, genomic, and tumor bulk RNA-seq analysis to find “aberrant” cells in normal human kidney whose gene expression profile matches that of malignant kidney tumors, such as Wilms tumors and renal cell carcinomas (RCCs) (Young et al. 2018). In their analysis, scRNA-seq datasets from the fetal (n = 2; normal kidneys), children (n = 3 for normal kidneys and n = 3 for Wilms tumors), and adult samples (n = 5 for normal kidneys, n = 1 for papillary RCCs, and n = 3 for clear cell RCCs) were generated and analyzed independently, and then the similarity between aberrant cells and tumors was ultimately identified from comparisons of the gene expression of individual cell clusters to the tumor bulk RNA-seq data and tumor markers. We thus tested whether RISC could reproduce the finding by simply performing an integrated analysis of the scRNA-seq data from normal and malignant kidneys.

Before integration, we reanalyzed the fetal, children and adult scRNA-seq datasets independently, and demonstrated that RISC correctly separated the cell types identified by Yang et al, based on the expression of the marker genes described in their paper (Fig. S4A ~ C), i.e., tumor cells (Wilms and RCC) and tumor-precursor nephrogenic rest (NR) cells were distinguished from cell types in the normal kidneys, including endothelium (EN), epithelial (EP), myofibroblast (MB), fibroblast (FB), cap mesenchyme (CM), ureteric bud (UB), primitive vesicle (PV), and Intermediate population (IP). When the fetal and children datasets were integrated, RISC largely removed the batch effects (Fig. S4D); and, more importantly, showed that the fetal PV and UB cells were embedded around a subset of Wilms tumor and NR cells (circled clusters in Fig. 5A), in agreement with the original finding that fetal UB and PV cells could be the origin of Wilms tumors (Young et al. 2018). Distinct from Seurat or Scran, (Fig. S4E), the RISC-integrated dataset showed that these cells were co-localized to the cluster FW5 and FW7, but segregated away from other normal kidney cell clusters.

**Figure 5.**
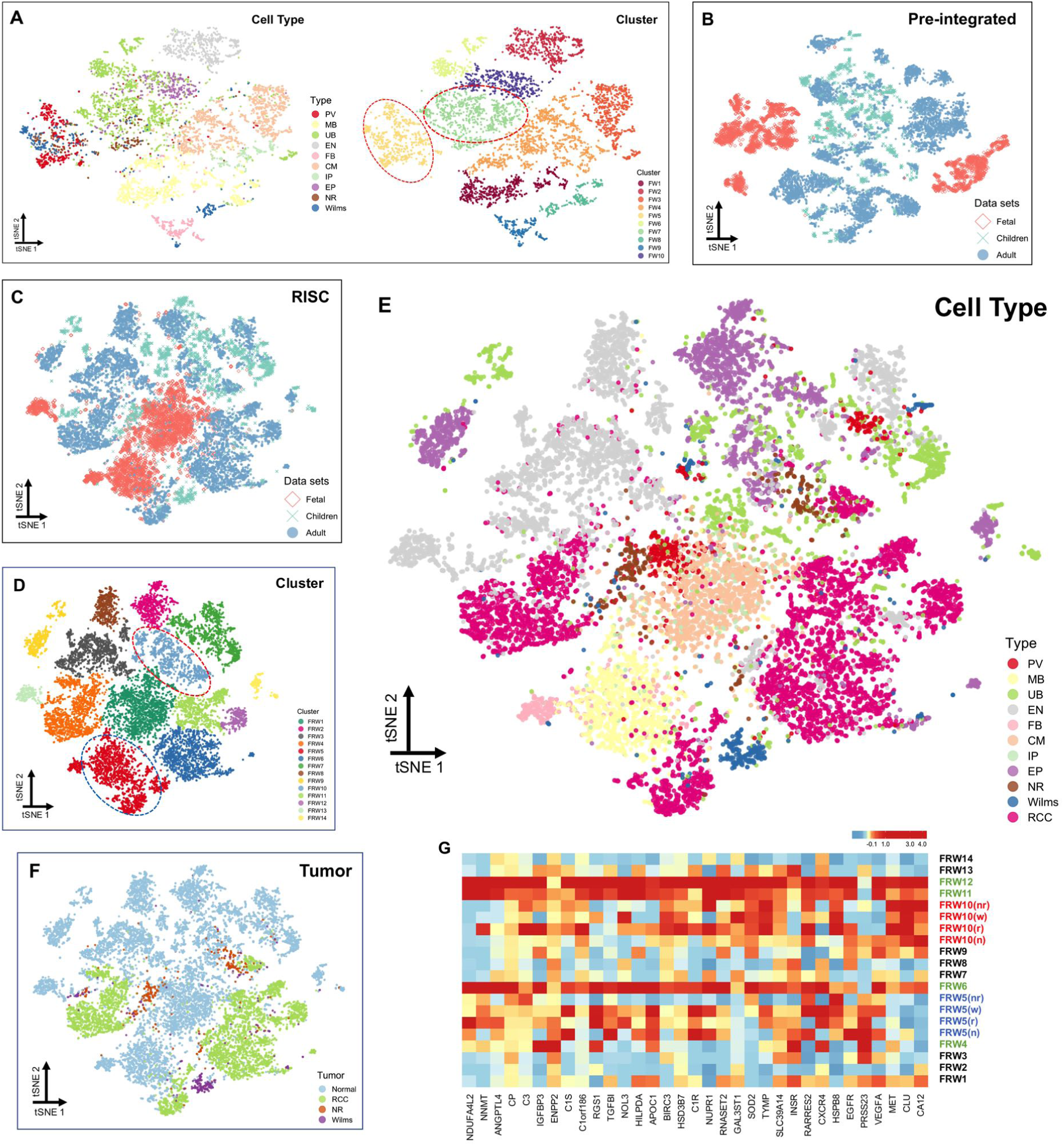
RISC integrated analysis of kidney normal cells and tumors. (A) T-SNE plots display cell types (left) and clusters (right) of the RISC-integrated data from the fetal and children datasets, including normal and tumor cells, colors on the left and right panels representing cell types and cell clusters, respectively. The red circles highlight the two clusters containing PV cells and Wilms tumors or UB and NR cells. (B ~ C) The pre-integrated (B) and RISC-integrated (C) datasets colored by their three sources, fetal, children and adult samples. (D) Clusters of the RISC-integrated data in (C), with the clusters PRW5 and PRW10 circled for containing both normal cells and tumors. (E) The cell types in the RISC-integrated data, consistent with Fig. S5A ~ C. (F) T-SNE plot displays the locations of the tumor cells from the original report (Young et al. 2018). (G) Heatmap displays expression patterns of 33 Wilms- or RCC-specific genes in kidney normal and tumor cells. The cluster FRW5 (navy for cluster label) and FRW10 (red for cluster label) are further divided into four subgroups: FRW5/10 (n) for normal cells, FRW5/10 (w) for Wilms tumors, FRW5/10 (r) for RCCs and FRW5/10 (nr) for NR cells. The green and black colors in cluster label represent tumor or normal-cell cluster, respectively.

When RISC was applied to integrate all kidney normal and tumor cells from fetal, children and adult samples, the batch separation was mitigated as more cells from the three sample groups became mixed (Fig. 5B ~ C). While normal kidney cells and tumor cells were generally separated, some normal cells were clustered together with tumor cells after integration (Fig. 5D ~ E). To help elucidate the identities of those cells, we projected the cell annotations by Yang et al onto the RISC-integrated and clustered data (Fig. 5F) (Young et al. 2018). The result indicated that two clusters contained both normal and tumor cells; FRW5 contained fetal MB cells, Wilms tumors and some RCCs, while the FRW10 cluster contained UB, EP, NR cells, and a different set of RCCs.

To further characterize these two cell clusters, we performed cluster marker analysis (Fig. S5F) and identified 1,020 and 192 genes highly expressed in the clusters FRW5 and FRW10, respectively, (Table S1, adjusted p-value < 0.05), including proto-oncogenes like KRT8 and OLFML3. The signature genes for the cluster FRW5 were enriched in oxidative phosphorylation (FDR = 2.6e-23), degradation of GLI2 by the proteasome (FDR = 6.9e-11), G2/M Transition (FDR = 1.2e-10) and other terms. The enriched pathways of the FRW10 cluster included apoptotic process (FDR = 5.1e-5) and others. We next searched for genes that were significantly differentially expressed (FDR < 0.05) between tumor clusters (FRW4, 6, 11 and 12) and normal-cell clusters (FRW1-3, 7-9, 13 and 14) (Fig. 5D, F) and were markers for at least one of the tumor clusters but not for the normal-cell clusters. Of the 33 genes meeting these criteria (Fig. 5G), many of them remarkably also showed high expression in the FRW5 and FRW10 clusters, including oncogene EGFR (Zandi et al. 2007) and genes linked to poor prognosis of patients with renal tumors, based on the data in the Human Protein Atlas database (Uhlen et al. 2017), such as TGFBI (Table S2). In short, the RISC-integrated data uncovers two small groups of normal kidney cells that exhibited high expression of several renal-cancer relevant markers and overall similar gene expression profiles to either Wilms tumors or RCCs, as indicated in the t-SNE plots.

## Discussion

Data integration is essential for extracting meaningful biological information across multiple scRNA-seq datasets, in order to decipher the cell-cell relationship across studies (Regev et al. 2017; Han et al. 2018). To facilitate robust data integration, we have developed RISC and evaluated its performance over multiple scRNA-seq datasets containing small to large portions of cell populations with matched gene expression profiles. In comparison to two currently available software, Seurat and Scran, RISC showed similar or better performance in merging cells with highly comparable cell compositions, but significant improvement in integrating datasets with subtle but biologically meaningful differences imbedded in otherwise homogenous datasets.

One advantage of RISC is its tendency to averting over integration when presented with datasets containing subtype difference, e.g., same cell types but at distinct functional states (Fig. 4). When more scRNA-seq studies begin to investigate cell functional difference beyond cell type identification, this feature can become very valuable. This feature can also be important for not to integrate unrelated datasets. To illustrate it, we incorporated a brain scRNA-seq dataset as a negative control (GSE103723) (Fan et al. 2018), and integrated it to the samples that we used for Figure 2, 3 and 4 separately. As Fig. S5A ~ S5C shown, cell integrating only happened to the same cell types (identified by cell-type marker genes used in Figure 2, 3 and 4) and brain cells were rarely mixed with non-brain cells. In practice, we expect users to analyze individual datasets and evaluate whether any of the cell clusters between two datasets share highly expressed gene markers before data integration, in order to ensure that the integrated result is biologically meaningful. In the future, we plan to develop a quantitative metric for evaluating the cell cluster similarity of two datasets and uses it as a guidance for integration.

The CCA, used in Seurat and Scran, is a classic statistic method for inferring signal information from cross-covariance matrices. However, the CCA has its limitation in handling matrices, in which the number of cells (matrix columns) is larger or similar to the number of genes (matrix rows). The regularization in CCA can solve this problem and has been developed for years, with several methods efficiently implementing the algorithm (Gonzalez et al. 2008; Witten et al. 2009), such as regularized CCA / ridge regression. These methods involve a penalizing parameter “lambda”, but how to determine the optimal value of lambda becomes a new challenge. In regularized CCA, one approach is to sample series of values for determining the optimal lambda that yields good fit error in CCA model. This requires a large number of repeated samplings and becomes extremely time consuming in scRNA-seq analysis. For instance, if we want to find the optimal lambda for two matrices, in the range of 0 to 1 with ten intervals, we have to perform CCA 100 times, each with a different lambda (Gonzalez et al. 2008). This is computationally infeasible for scRNA-seq integration with tens of thousands of cells in each dataset. Therefore, the lack of efficient method for identifying optimal lambda restricts the application of regularized CCA in scRNA-seq analysis. In RISC, instead of estimating this lambda, the PCR model selects the PCs based on dimension reduction, the process regularizes the matrices and generates the unique singular vectors at the first step of scRNA-seq data analysis. In addition, the PCs from dimension reduction preserves the meaningful biological information of each dataset, so the mathematical framework of PCR model allows it to avoid any noise that would not be used in the data analysis of individual datasets.

At last, we compared the running time of RISC, Seurat and Scran, from integrating individual datasets to generation of the final integrated expression matrix, and found that interestingly, the running time of Scran was completely dependent on how many cell pairs were identified from the MNN. The running time of RISC is significantly less than that of Seurat in all samples tested in the current study (Fig. S6). Although other well-known regularized models, such as lasso and partial least square (PLS) regression, also prevent overfitting and are suitable for gene matrices, a large number of time-consuming calculations would limit the practical use of these algorithms in most scRNA-seq analysis. Nevertheless, RISC provides an alternative PLS model by using SIMPLS algorithm, but the complicated computation looping reduces its performance.

## Data and Methods

### Description of the scRNA-seq datasets

#### PBMCs datasets

Two single-cell gene expression count matrices were generated from peripheral blood mononuclear cells of systemic lupus erythematosus (SLE) and Rheumatoid arthritis (RA) patient donors by Kang et al. (GSE96583) (Kang et al. 2017), with IFN-beta treatment in 6 hours (stimulation) or without treatment (control). The cells were isolated by Ficoll and prepared by the 10X genomics protocol. After cell filtration by RISC, 6,573 cells were valid in control (GSM2560248) and 7,466 cells in stimulation samples, with ~700 minimum UMIs/counts and ~500 expressed genes in each cell. Totally, 6 cell types were identified by marker genes, including B, NK, T, CD14 monocytes, CD16 monocytes and DC cells; among them, T cells were further divided into four sub-populations, CD4 Naïve T, CD8 T, activated T and CD4 Memory T cells (Kang et al. 2017; Butler et al. 2018).

#### Human and mouse pancreatic-islet single-cell datasets

The single-cell datasets were derived from pancreatic islets of four human donors and two mouse strains by Veres A. et al. (GSE84133) (Baron et al. 2016), using either inDrop workflow, microfluidic droplet platforms, or the CEL-seq protocol. The human cells were from two males and two females, while two mice were the ICR and C57BL/6 strains. After cell filtration by RISC, we got 8,100 valid cells for human and 1,781 cells for mice, with ~11,000 genes expressed in both datasets. Using the cell type annotation from the original report (Baron et al. 2016), we identified ten cell populations, including acinar, alpha, beta, gamma, delta, ductal, endothelial, stellate and macrophage cells.

#### Human pancreas single-cell datasets

Three pancreas single-cell datasets were obtained from either the ArrayExpress (E-MTAB-5061) (Segerstolpe et al. 2016) or the GEO (GSE81076 and GSE85241) (Grun et al. 2016; Muraro et al. 2016). The E-MTAB-5061 was prepared by the Smart- seq2 protocol. GSE81076 was collected by the CEL-seq and GSE85241 by the CEL-seq2 protocol. The E-MTAB-5061 data contained ten donors, six healthy and four T2D (type 2 diabetes) cadaveric patients. The single cells in GSE81076 were derived from organ donors with or without T2D, and four healthy human donors provided single cells for the GSE85241 data. After preprocessing, 680 cells passed quality control in E-MTAB-5061 with the minimum of 1,000 expressed genes and 1,000 UMIs/counts per cell, 2,241 single cells for GSE81076 with the cutoff of 1,000 expressed genes and 3,000 UMIs/counts, and 1,140 valid cells in GSE85241 with more than 1,000 expressed genes and 1,000 UMIs/counts. The integrated analyses of RISC, Seurat and Scran all identified four cell sub-populations, marked by specific genes (GCG, INS, PPY and SST for alpha (α), beta (β), gamma (γ) and delta (δ) cells, respectively) (Grun et al. 2016; Muraro et al. 2016; Segerstolpe et al. 2016).

#### HSPC datasets

Two single-cell datasets of hematopoietic lineage were downloaded from the GEO databases (GSE81682 and GSE72857), and contributed by Nestorowa et al. (Nestorowa et al. 2016) and Amit et al. (Paul et al. 2015), respectively. In the GSE81682, single cells were captured from hematopoietic stem and progenitor populations and followed by Smart-seq2 protocol for library preparation. The cells in GSE72857 were isolated from Bone marrow Lin- cKit+ Sca1- myeloid progenitor cells, and processed with the MARS-seq protocol. After filtering, 803 cells in GSE81682 and 2,679 cells in GSE72857 passed our quality control, with ~8,400 expressed genes in the integrated data. According to the original reports (Paul et al. 2015; Nestorowa et al. 2016), Gata1, Irf8 and Mpo were used as markers for erythrocyte progenitors, monocyte progenitors, and neutrophil progenitors, respectively.

#### T cell datasets from breast cancers

Three datasets (BC09, BC10 and BC11) for single T cells were derived from breast cancer patients by Azizi et al. (GSE114727) (Azizi et al. 2018), and all cells were CD3+ leukocytes and sequenced in paired V(D)j platform by the 10X genomics workflow. Both BC09 and BC11 had two technical replicates, but in this study, we only used one replicate (GSM3148575 for BC09 and GSM3148578 for BC11); by contrast, BC10 only had one replicate (GSM3148577). We first filtered the cells based on the authors’ single cell T cell receptor sequencing annotation (Azizi et al. 2018). Then, after preprocessing by RISC, 6,550 single cells passed the quality control for BC09, 4,593 for BC10 and 4,982 for BC11, with ~12,500 expressed genes across three datasets. These cells were divided into three major types were, including CD8+, CD4+ and regulatory T cells (Treg), according to T cell activation signature genes (CD8A, CD4 and FOXP3) (Azizi et al. 2018). A second T cell scRNA-seq dataset was also from breast cancer patients by Savas et al. (GSE110686) (Savas et al. 2018); all cells were CD3+ and isolated by fluorescence-activated cell sorting (FACS) and sequenced by by the 10X genomics workflow. After preprocessing by RISC, 5,990 single cells passed the quality control, with ~12,4000 expressed genes.

#### Kidney normal and tumor datasets

The scRNA-seq datasets of human kidney normal and tumor cells were previously generated from fetal, children and adult patients or donors (Young et al. 2018), including ~72,000 single-cell transcriptomes in total. Filtering out the cells labeled as “low quality”, immune cells, and normal proximal tubule cells not related to tumors (as defined by the original report), we kept ~18,000 single-cell transcriptomes for this study. These single cells were derived from the fetal (n = 2 for normal kidneys), the children (n = 3 for normal kidneys and n = 3 for Wilms), and the adult (n = 5 for normal kidneys, n = 1 for papillary RCCs, and n = 3 for clear cell RCCs) samples. After data preprocessing, 17,067 cells passed the quality control of RISC, with ~16,000 expressed genes in common. As the original report provided the cell-type markers and labels the tumor-status of each cell, here we used these information as they were in the supplemental “Table” file of the original report (Young et al. 2018).

#### Brain negative control dataset

The dataset was previously derived from single embryonic cortex cells of 22-23 weeks human embryos, using the Smart-seq2 protocol by Fan et al. (GSE103723) (Fan et al. 2018). After data preprocessing, 4,408 cells with ~20,000 expressed genes were kept for our analysis.

#### Filtering and processing scRNA-seq data

We filtered all scRNA-seq datasets using standard normal distribution to remove cells with extremely low/high UMIs or low/high number of expressed genes. We also discarded the genes only expressed in few cells, according to distribution analysis. The same filtered datasets were used as inputs for RISC, Seurat and Scran.

To remove the technical batch effect of sequencing depth, we normalized the gene expression values of individual datasets by size factors that were equal to the total number of transcripts in each cell (i) divided by median value of all cells.

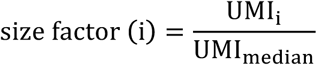

We further scaled the gene expression values of all cells in each dataset by Poisson scaling (root-mean-squaring), and centered gene expression magnitude to the same level with empirical mean equal to 0. The root-mean-squaring scaled matrices preserve gene expression signals used for data integration.

Identify highly variable genes by Quasi-Poisson model. The default method in RISC utilizes three criteria to pick up highly variable genes. First, coefficient of variation is calculated for each gene, given by

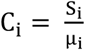

where S_i_ and μ_i_ denote standard devotion and mean value, respectively, for gene i (i ∈ 1, 2, …, n), and the genes with C_i_ > 0.5 are selected. To control for the relationship between S_i_ and μ_i_, Quasi-Poisson regression is used to further filter genes with over-dispersion C caused by small μ (Ver Hoef and Boveng 2007). Specifically, genes are put into a set of bins (e.g. 20 bins) based on their expression level. For genes in a bin (b ∈ 1, 2, …, 20), Quasi-Poisson regression is used to predict 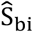 by μ_bi_ as

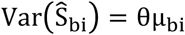

where θ for the Quasi-Poisson over-dispersion parameter. The predicted 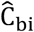 is calculated by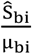 and the corresponding ratio between the observed Ci and the predicted 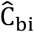 is given by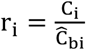 for each gene. The genes with r > 1 is considered as highly variable for further subsequent analysis. We limit the number of the highly variable genes < 1,500 by ranking r_i_, with z-score test.

Besides the Quasi-Poisson regression model, RISC also provides an alternative method, named “residuals”, to choose the highly variable genes, and this method is similar to what is used in the Seurat package (Butler et al. 2018).

#### Data integration by PCR

The principal PCR model is designed to identify a covariance hyper- plane between two matrices. To illustrate the idea, we consider two datasets (two matrices of gene expression values). The extension to multiple datasets is straightforward.

#### PCR model in two datasets

The core of PCR is based on an ordinary least square linear regression. Let two gene-expression-value matrices be a reference n×p matrix X_n×p_ with n rows of genes and p columns of cells and an n×q matrix Y_n×q_. PCR provides the estimated regression coefficients between the two matrices (also shown in Fig. 1B) given by

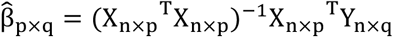

in which p ≥ q due to the reference dataset is selected to have the larger cell number.

PCR model is based on the principal component (PCs) of the matrix X_n×p_. Thus, under the condition that

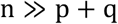

(the condition n ≅ p + q or n ≤ max (p, q) will be discussed in “Regularized procedure in PCR model” section), the matrix can be decomposed using SVD. Then, after removing the uninformative components (n - p), the matrix is given by

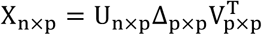

where Δ_*p*×*p*_ denotes the singular values, and *U*_*n*×*p*_ and *V*_*q*×*q*_ represent the left- and right-singular vectors of the matrix *X*_*n*×*p*_, respectively. Hence,

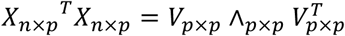

where ^_*p*×*p*_ denotes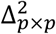.

When all PCs are used, the regression coefficient estimates of PCR are equivalent to the product of two matrices. Of note, PCR only uses top PCs, which is different with the principal CCA that is equivalent to an approach using all PCs. Let

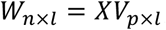

where *V*_*p*×*l*_ is the corresponding submatrix of *V*or *l* ∈ 1, 2, …, *p*. Therefore, we have

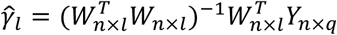

and

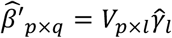

In briefly, the coefficient matrix 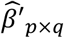between *X*_*n*×*p*_ and *Y*_*n*×*q*_ is estimated and dependent on the first *l* PCs in PCR model.

#### Regularized procedure in PCR model

In most cases, the dimension reduction and data integration of scRNA-seq data are based on the highly variable genes (usually *n* ≤ 1,500). Since the cell number of scRNA-seq data are often thousands and more, the matrix *X* _*n×p*_ and *Y*_*n×p*_ are most likely in the condition

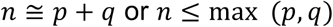

Thus, unique SVD may not be obtained from either *X* _*n×p*_ or *Y*_*n×p*_ in real scRNA-seq data. This issue cannot be handled by the principal CCA, and needs to be corrected by the regularized CCA (Gonzalez et al. 2008). In contrast, the PCR model itself undergoes the regularized solution. The PCR model utilizes the first *l*-column PCs of *X* _*n×p*_, and in practice *l* ≪ *n*, so the *V*_*p*×*l*_ and *W*_*n*×*l*_ will be unique in PCR model. Our challenging is to find the optimal *l* that can preserve most important biological information of datasets, and to solve the constrained minimization problem of the rest (*q* – *l*) columns of PCs through

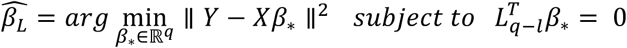

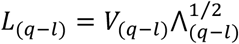

However, we know the optimal first *l* -column PCs is equal or approximate to the maximum number of PCs used for dimension reduction in each dataset, so the PCR model forms a regularized procedure by itself when we insure *l* ≪ *n*.

#### Alignment of gene expression values

For the downstream scRNA-seq analysis, RISC aligns gene expression values across datasets by two options. Primarily, according to PCR model, one can predict the target-matrix values based on the reference matrix *X*_*n*×*q*_ directly, i.e. in data integration of two datasets, the predicted 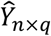is given by

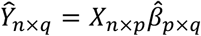

where 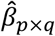denoting regression coefficient matrix. Then, the expression range of each gene is adjusted. Since the minimum of normalized counts in gene expression matrix is zero, 99.5% quantiles of individual genes denote the expression ranges of the genes in all cells.

Instead of simply balancing every gene to have the same expression range, we first get a set of range vectors for all shared expressed genes between two datasets.

Let

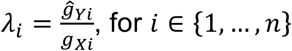

denote range vectors for all the genes, where *g*_*Xi*_ represents expression range of gene *g*_*i*_ in the reference matrix *X*_*n*×*p*_ and 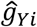for expression range of gene *g*_*i*_ in the predicted matrix 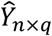. Based on our kernel assumption, a large number of genes is not differentially expressed across datasets, we thus utilize log-linked linear regression model with Gaussian distribution to predict the empirical confidence intervals of *λ_i_*,

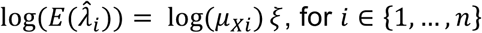

where *μ_Xi_* denoting average expression value of gene *g*_*i*_ in the reference matrix *X*_*n*×*p*,_ and ξfor a linear coefficient based on *λ_i_* byμ*_Xi_*. The genes are considered to be in different expression ranges across datasets, with 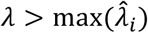 or 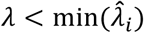 for *i* ∈ 1, …, *n*. For multiple datasets, the processes are repeated between the reference and the rest datasets respectively, then the values of all the matrices are modified according to the reference dataset and made comparable for the downstream analysis.

An alternative method in RISC is to skip the step of gene-expression-value prediction according to PCR model. It is still based on the kernel assumption that most genes are not differentially expressed across datasets. Thus, we directly adjust the expression ranges of each gene. Since PCR-prediction will transform the variance structures of multiple datasets to be similar to the variance structure of the reference dataset. This alternative method can optimally reserve the original variance structures of individual datasets.

#### Cell clustering in RISC

Three cell clustering methods are provided in RISC. The first is based on the k-nearest neighbor (KNNs) algorithm, the second is by Gaussian Mixture Models (GMMs), and the last extends and improves from density peak clustering algorithm. These clustering methods were previously described in R packages “densityClust” (Rodriguez and Laio 2014) and “ClusterR” (Maechler 2018). The default method (used in this report) is the “density-based clustering”.

#### Running of Seurat and Scran

We used the default parameters to run Seurat (v2.3.4) and Scran (v1.8.0) in all analyses. The normalization and scaling of gene expression values were performed by Seurat or Scran themselves. The top variable genes for running Scran were taken from the Seurat, as Scran does not identify top variable genes and both software utilize the algorithm CCA. We should point out that the default t-SNE function in RISC, Seurat and Scran (Scran calculating t-SNE using the “scater” package) calls the same function in the “Rtsne” package (van der Maaten 2014; McCarthy et al. 2017; Butler et al. 2018; Haghverdi et al. 2018), so the t-SNE plots from the three software are directly comparable.

## Data and code availability

All scRNA-seq datasets in this study have been published by other research groups. The RISC is prepared as a R package and will be released as a R CRAN package for free usage.

## Supplemental Figures

**Figure S1.**
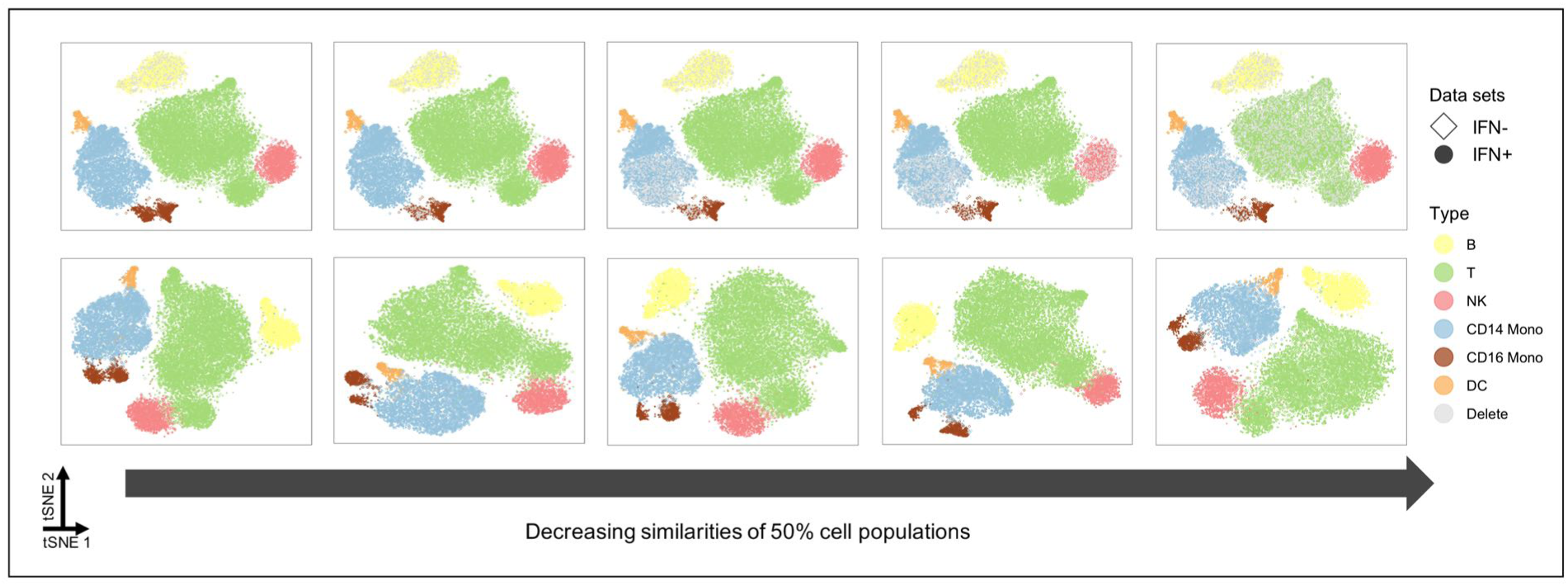
The performance of RISC in data integration with 50% reducing cell-population similarities. The upper plots display the “IFN+” data subsets with reduction of cell-population similarity to the “IFN-” data (from left to right), gray color marking the 50% cells removed. The lower plots show cell embedding of the integrated dataset between the “IFN-” data (full cell populations) and the individual “IFN+” subsets. The colors for cell populations correspond to Fig. 2C.

**Figure S2.**
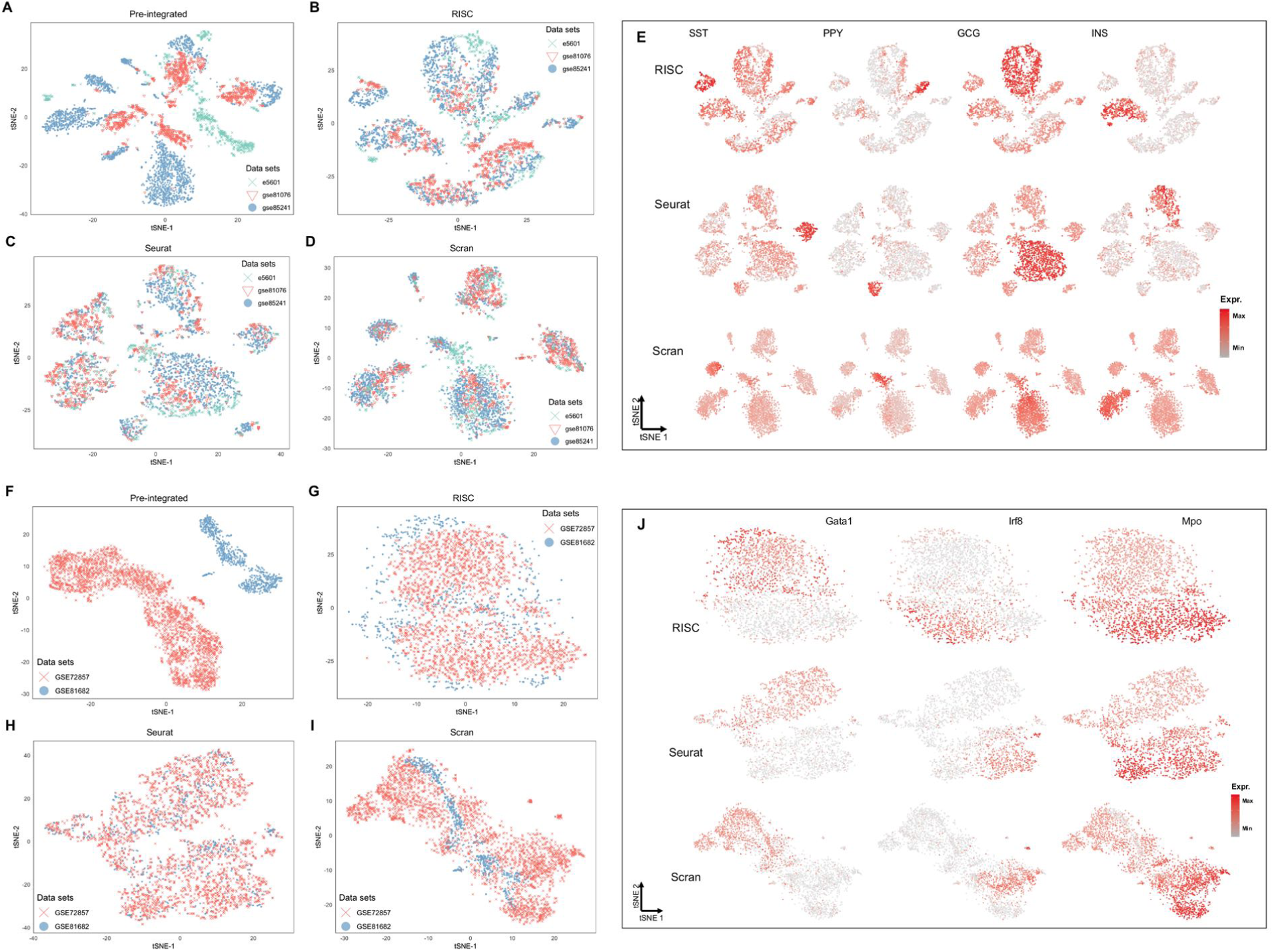
Integration of data from different studies and platforms. (A ~ D) The t-SNE plots show the cell embedding of the pre-integrated datasets (A) from three pancreas scRNA-seq data, and the integrated datasets by RISC (B), Seurat (C) and Scran (D). (E) The plots display expression patterns of cell-type marker genes. (F ~ I) The t-SNE plots indicate the cell embedding of the pre-integrated datasets (F) of the hematopoietic lineages from different platforms, and the integrated datasets by RISC (G), Seurat (H) and Scran (I). (J) The expression patterns of the known cell-type markers.

**Figure S3.**
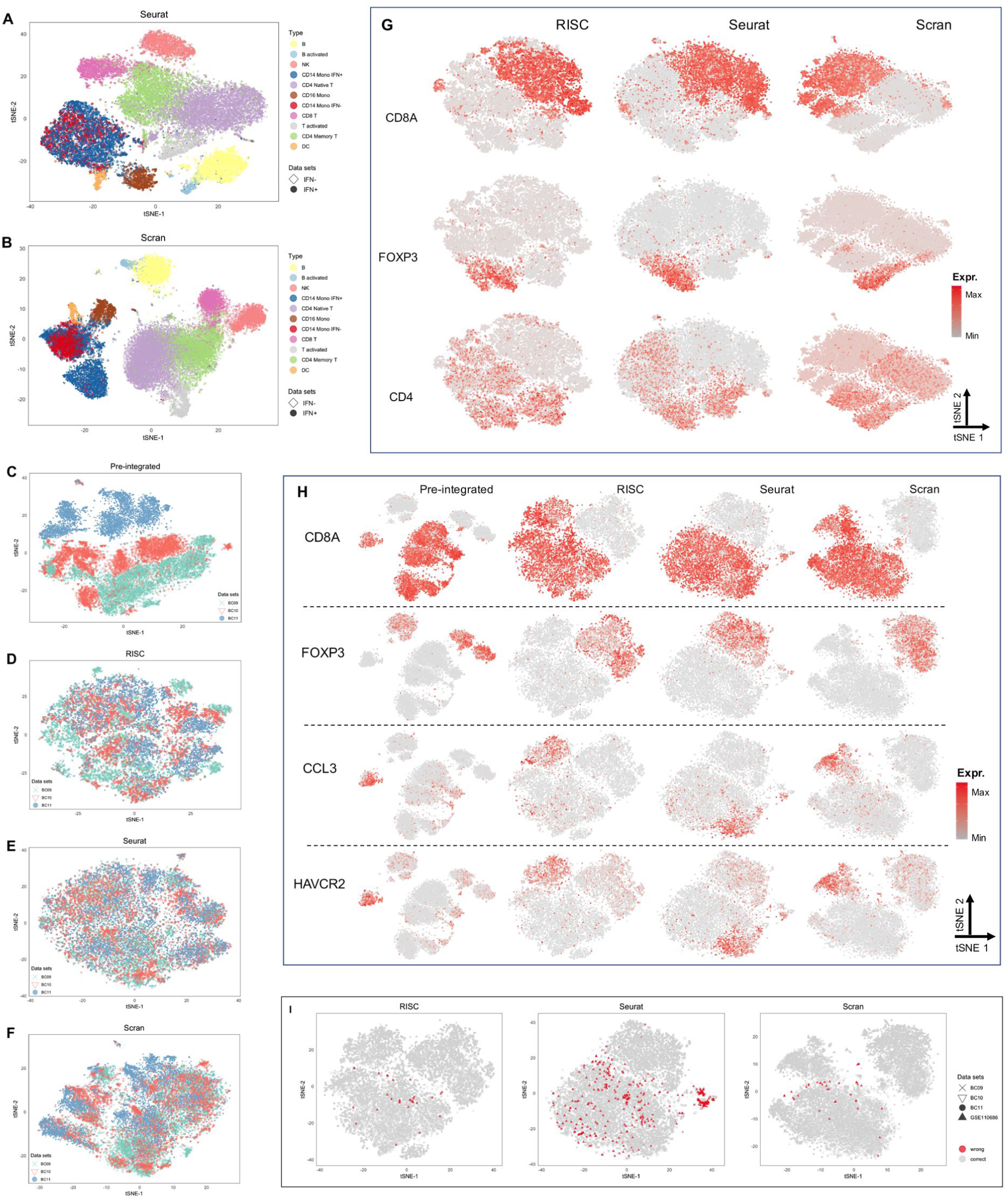
Integration of full T-cell datasets. (A ~ B) The cell types of the Seurat-integrated (A) and Scran-integrated (B) data from the two PBMC datasets in Fig. 2, the colors for cell types corresponding to the colors in Fig. 2D. (C ~ F) Comparison of the data before (C) and after integration by RISC (D), Seurat (E) and Scran (F). The three colors indicate T cells from three breast cancer patients (BC09, BC10 and BC11). (G) Expression patterns of marker genes (CD4, CD8A and FOXP3) for three distinct T cell types. (H) Expression patterns of marker genes (CD8A, FOXP3, CCL3 and HAVCR2) for CD8+ Trm-, CD4+ Treg and CD8+ Trm+ cells. (H) T-SNE plots display the CD8+ Trm cells that were integrated incorrectly to non-Trim cells.

**Figure S4.**
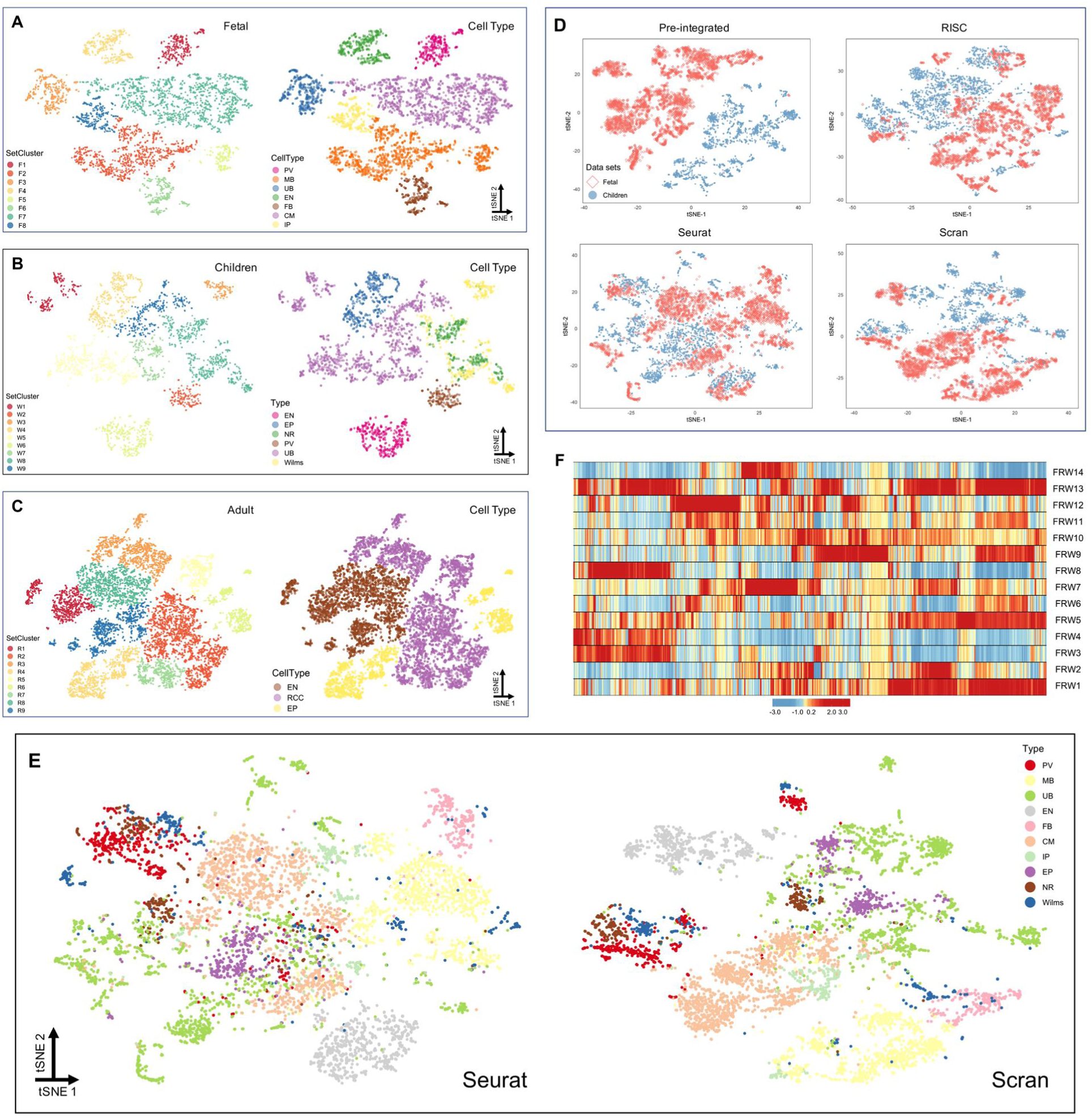
Application of RISC to normal and malignant kidney cells. (A ~ C) The t-SNE plots show cell clustering (left) and cell types (right) of the fetal (A), children (B) and adult (C) datasets by RISC. (D) Comparison of the fetal and children datasets before and after integration by RISC, Seurat or Scran, with cells colored by their sources. (E) The t-SNE plots highlight the cell types in the integrated datasets (D) by Seurat or Scran, respectively. (F) The heat-map shows expression patterns of the top cluster markers in the RISC-integrated data, from the full three datasets consistent to Fig. 5D.

**Figure S5.**
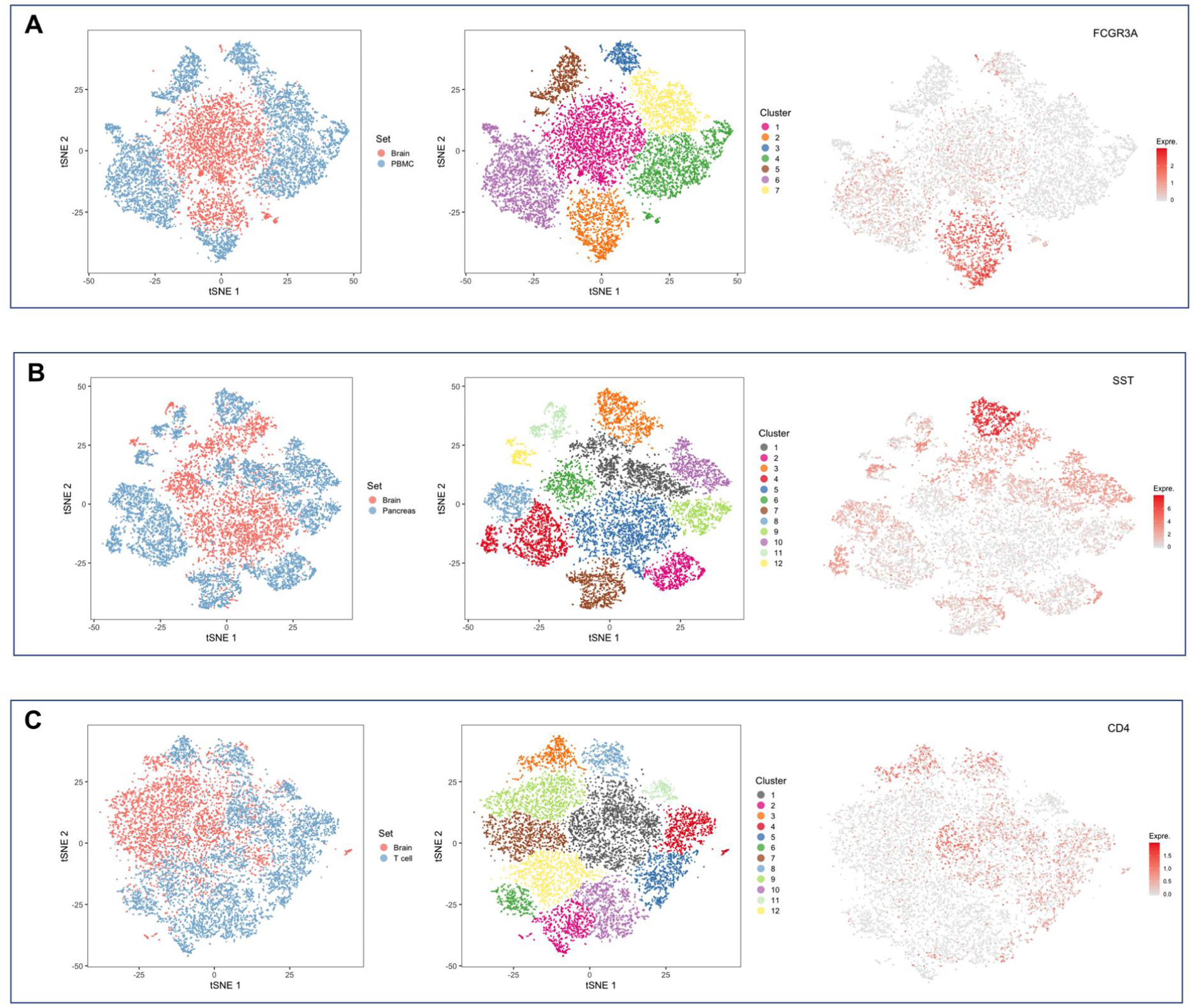
No integration of brain to non-brain cells. (A ~ C) The combination of single-cell profiles of human brain to PBMC (A), pancreas (B), or T cell (C) datasets, respectively. In each panel, middle t-SNE plot shows clusters of the integrated data and right plot for expression pattern of marker gene in the integrated data.

**Figure S6.**
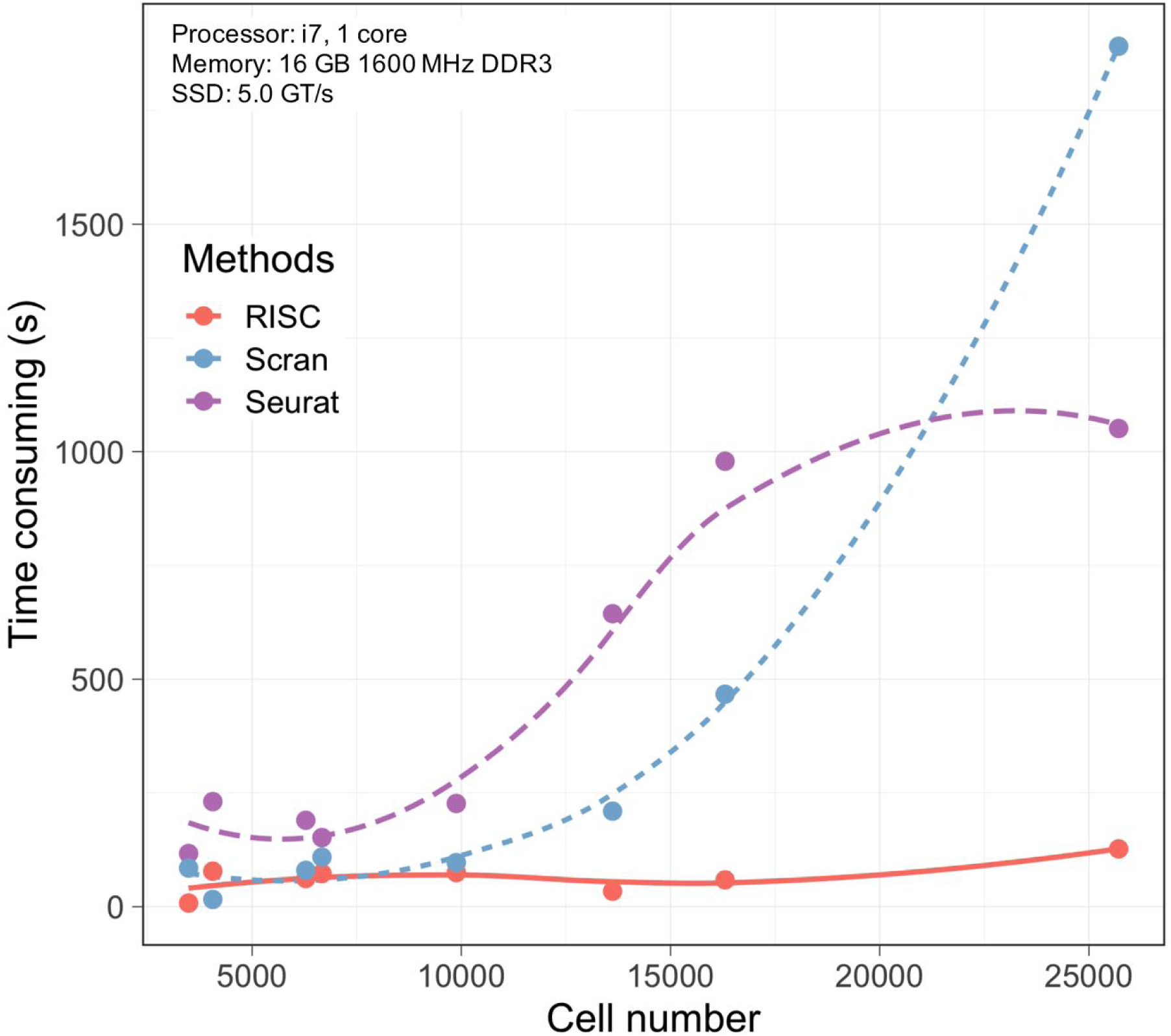
The runtime of RISC, Seurat and Scran in data integration. Each point in the plot represents the running time for a specific data integration described in this study.

## Supplemental Tables

**Table S1.** Cluster markers for the integrated liver scRNA-seq data

**Table S2.** Tumor specifically expressed genes

